# Desmoplakin Haploinsu/iciency Underlies Cell-Cell Adhesion Failure in DSP Cardiomyopathy and is Rescued by Transcriptional Activation

**DOI:** 10.1101/2025.06.01.657304

**Authors:** Eric D. Smith, Karen Jin, Brianna Ferguson, Yao-Chang Tsan, Samuel J DePalma, Joshua Meisner, Aaron Renberg, Kenneth Bedi, Sabrina Friedline, Kenneth B. Margulies, Brendon M. Baker, Adam S. Helms

**Affiliations:** Department of Internal Medicine, Division of Cardiovascular Medicine, University of Michigan, Ann Arbor, MI, USA; Department of Biomedical Engineering, University of Michigan, Ann Arbor, MI, USA; Cardiovascular Institute, Perelman School of Medicine, University of Pennsylvania, Philadelphia, PA, USA

## Abstract

**Background:** Truncating variants in desmoplakin (*DSP*tv), are a leading cause of arrhythmogenic cardiomyopathy (ACM), often presenting with early fibrosis and arrhythmias disproportionate to systolic dysfunction. DSP is critical for cardiac mechanical integrity, linking desmosomes to the cytoskeleton to withstand contractile forces. While loss-of-function is implicated, direct evidence, both for DSP haploinsufficiency in human hearts and for the impact of mechanical stress on cardiomyocyte adhesion, has been limited, leaving the pathogenic mechanism unclear.

**Methods:** We analyzed explanted human heart tissue from patients with *DSP*tv (N=3), titin truncating variants (*TTN*tv, N=5), and controls (N=5) using RNA-sequencing and mass spectrometry. We generated human induced pluripotent stem cell-derived cardiomyocytes (iPSC-CMs) harboring patient-derived or CRISPR-Cas9 engineered *DSP*tv to model a range of DSP expression levels. Using a 2D cardiac muscle bundle (CMB) platform enabling live visualization of cell junctions, we developed an assay to assess cell-cell adhesion upon heightened contractile stress in response to the contractile agonist endothelin-1. CRISPR-interference (CRISPRi) was used to confirm the role of DSP loss, and CRISPR-activation (CRISPRa) was tested for therapeutic rescue.

**Results:** Compared to both control and *TTN*tv hearts, *DSP*tv human hearts exhibited reduced *DSP* at both the mRNA and protein level, as well as broadly disrupted desmosomal stoichiometry. Transcriptomic and proteomic analyses implicated cell adhesion, extracellular matrix, and inflammatory pathways. iPSC-CM models recapitulated DSP haploinsufficiency and desmosomal disruption. *DSP*tv CMBs showed normal baseline contractile function. However, they displayed marked cell-cell adhesion failure with contractile stress (75% failure vs. 8% in controls, p<0.001). Adhesion failure was prevented by the myosin inhibitor, mavacamten. CRISPRi-mediated DSP knockdown replicated this susceptibility to adhesion failure.

Conversely, CRISPRa robustly increased DSP expression and rescued cell-cell adhesion failure in *DSP*tv CMBs (9% failure post-CRISPRa, p<0.001 vs. un-treated). Rescue occurred even when only the DSPII isoform was upregulated in a model with biallelic *DSP* transcript 1 loss of function.

**Conclusions:** DSP haploinsufficiency is the major cause of DSP cardiomyopathy with a primary consequence of conferring vulnerability to cardiomyocyte cell-cell adhesion failure under heightened contractile stress. Transcriptional activation of *DSP* reverses this defect in preclinical models, establishing proof-of-concept for a potential therapeutic strategy in DSP cardiomyopathy.

## Introduction

Variants in the gene desmoplakin (*DSP*) are the most common cause of the dilated cardiomyopathy (DCM) subcategory, left-dominant arrhythmogenic cardiomyopathy (ACM).^1, 2^ *DSP* variants account for 10% of all genetically defined DCM/ACM.^2^ Clinical features that distinguish DSP cardiomyopathy include early onset fibrosis in the subepicardial muscle layer and ventricular arrhythmias out of proportion to the degree of systolic dysfunction. Most reported pathogenic *DSP* variants are truncating variants (*DSP*tv), suggesting a loss of function mechanism. However, DSP haploinsufficiency has not been rigorously demonstrated in human heart tissue from patients with *DSP*tv.

In the heart, DSP links desmosome complexes via plaque proteins (plakoglobin and plakophilin-2) to the intermediate filament protein, desmin, which forms a structural scaffold that connects to the Z-disks of myofilaments.^3^ As such, DSP transduces force from the contractile machinery to intercalated disks and must be able to withstand cyclic tensile loading during cardiac contraction. DSP’s role in cellular resilience during tensile loading has been demonstrated in epidermal keratinocytes, in which DSP serves a similar structural role.^4–6^ Investigation of tensile loading stress on cell-cell adhesive function in DSP cardiomyopathy models has been limited due to the challenge of modeling and measuring cardiomyocyte cell junctions under physiologic contractile conditions. At the tissue level, *DSP* variants appear to result in reduced contractile function under higher preload, but the mechanism of this observation has not been clearly defined.^7–9^ Several additional hypotheses have been explored. *DSP* variants may disrupt functions of the desmosome as a signaling hub, altering regulation of Wnt/β-catenin, Hippo, ERK/p38MAPK, and gap junctions.^10–15^ Complete knock-out of DSP in engineered heart tissues was shown to cause a damage associated molecular pattern, including increased NF-κβ expression, similar to findings from a desmoglein-2 homozygous knock-in mouse model of ACM.^8, 16^ The extent to which these various pathologies could be downstream of a primary deficit in mechanical function and cardiomyocyte cell-cell adhesion is unknown.

Here, we definitively establish DSP haploinsufficiency in explanted human heart tissue from patients with DSP cardiomyopathy compared to both titin (*TTN*) cardiomyopathy (with a similar amount of ventricular remodeling) and control hearts. We then generated human induced pluripotent stem cell-derived cardiomyocyte (iPSC-CM) lines that exhibit a range of DSP haploinsufficiency, modeling the extent of DSP haploinsufficiency present in human hearts.

Leveraging a 2D cardiac muscle bundle approach that enables live cell monitoring of cell junctions in physiologically contracting cardiac tissues, we developed an assay that clearly identifies cell-cell adhesion failure as a primary disease mechanism in DSP models. We then demonstrate mechanistically that DSP loss of function drives cell adhesion dysfunction using CRISPR-Cas9 interference (CRISPRi). Finally, we show that CRISPR-Cas9 activation (CRISPRa) is capable of rescuing DSP expression and restoring normal cell adhesion function, despite a genetic background of severe DSP haploinsufficiency. These studies show proof-of-concept that targeting *DSP* transcription could be a viable therapeutic intervention for DSP cardiomyopathy arising from loss of function variants.

## Methods

### Human Heart Tissue Processing

Patients with *DSP*tv and *TTN* truncating variants (*TTN*tv) were identified from explanted heart tissue biobanks at the University of Michigan and University of Pennsylvania. The tissue biobanks were approved by the Institutional Review Board (IRB) of each institution, and subjects gave informed consent for research use of explanted tissue. Ventricular myocardial tissue was snap frozen in liquid N_2_ at the time of collection.

### Cell Culture

The use of iPSCs for this project was approved by the University of Michigan Human Pluripotent Stem Cell Research Oversight Committee. Control iPSCs were obtained from Coriell (WTC-11, GM25256), the Allen Institute (DSP-GFP, AICS-0017 cl.65), and Vanderbilt (CF-3).^17^ Donors of cells previously gave informed consent prior to inclusion in the respective repositories above, and their identities are anonymous. IPSCs were cultured in mTeSR Plus (StemCell Technologies) and verified free of mycoplasma using MycoAlert^TM^ (Lonza). Cardiac differentiations were performed using Wnt modulation with modifications as previously described.^18, 19^ We achieved optimal differentiations using RPMI 1640 plus B27 supplement without insulin during day 0-2 (CHIR 5-6 µM during first 24 hours, LC Laboratories; plus Activin A (R&D, #338-AC-010)), then CDM3 media (without B27) with IWP4 5 µM (Stemgent) and retinol inhibitor (BMS 453, Cayman Chemical, 1 µM) on day 3-4 to minimize atrial lineage differentiation.^18^ hPSC-CMs were maintained in CDM3 until purification in CDM3-lactate using glucose-deprived and glutamine-deprived, lactate-containing media for 4 days.^19^ Purified hPSC-CMs were replated as monolayers (400,000 cells/cm^2^) for 8 additional days (until day 24) on stiff PDMS membranes (Specialty Manufacturing, Inc.) with growth factor reduced Matrigel (Corning) in CDM3-lactate media (4 days) and then oxidation phosphorylation promoting maturation media as previously described (4 days).^18, 19^ At day 24, hPSC-CM batches with >95% cardiomyocyte purity were either collected for RNA or protein extraction, or they were replated onto cardiac muscle bundle substrates for functional analyses.

### Gene Editing

CRISPR-Cas9 editing was used to create truncating variants targeting only *DSP* transcript 1 by targeting the transcript 1-specific portion of exon 23. CRISPOR 4.98 was used to identify highly active gRNA target sites with low predicted off-target recognition. Two gRNAs were selected, and gene editing was performed as previously described (**Supplementary Table 1**).^20^ The DSP-GFP iPSC line from Allen Institute (AICS-0017 cl.65) was used for targeting. Single cell derived clones were expanded and genotyped by PCR and Sanger sequencing to identify clones containing either single allelic truncating or biallelic truncating variants. The results of CRISPR-Cas9 editing and targeting guides are shown in **Supplemental Figure 1**.

### Mass Spectrometry

Mass spectrometry was performed by the Mass Spectrometry Core Laboratory at the University of Texas Health Science Center at San Antonio. For human heart tissue, 2-3 mg of each thawed heart tissue was placed in a MicroTube fitted with a MicroPestle (Pressure BioSciences, Inc.), and 30 µl of homogenization buffer [10% SDS in 50 mM triethylammonium bicarbonate (TEAB, Thermo Fisher Scientific) containing protease/phosphatase inhibitors (Halt, Thermo Fisher Scientific) and nuclease (Universal Nuclease, Pierce/Thermo Fisher Scientific)] was added. The tissue samples were homogenized in a Barocycler (Pressure BioSciences, Inc.) for 60 cycles at 35 °C and then the MicroTubes were centrifuged at 21000x g for 10 minutes. For iPSC-CM samples, 1e6 iPSC-CMs were pelleted for each sample and homogenized in the same buffers but without requiring the Barocycler step. The supernatants were removed and stored at −80 °C until processed for mass spectrometry analysis. Aliquots corresponding to 100 µg protein (EZQ™ Protein Quantitation Kit; Thermo Fisher Scientific) were reduced with tris(2-carboxyethyl)phosphine hydrochloride (TCEP), alkylated in the dark with iodoacetamide and applied to S-Traps (mini; Protifi) for tryptic digestion (sequencing grade; Promega) in 50 mM TEAB. Peptides were eluted from the S-Traps with 0.2% formic acid in 50% aqueous acetonitrile and quantified using Pierce™ Quantitative Fluorometric Peptide Assay (Thermo Fisher Scientific). Data-independent acquisition mass spectrometry was conducted on an Orbitrap Fusion Lumos mass spectrometer (Thermo Fisher Scientific). On-line HPLC separation was accomplished with an RSLC NANO HPLC system (Thermo Fisher Scientific/Dyonex): column, PicoFrit™ (New Objective; 75 μm i.d.) packed to 15 cm with C18 adsorbent (Vydac; 218MS 5 μm, 300 Å); mobile phase A, 0.5% acetic acid (HAc)/0.005% trifluoroacetic acid (TFA) in water; mobile phase B, 90% acetonitrile/0.5% HAc/0.005% TFA/9.5% water; gradient 3 to 42% B in 120 min; flow rate, 0.4 μl/min. A pool was made of all of the samples, and 2-µg peptide aliquots were analyzed using gas-phase fractionation (three separate injections, together covering m/z 400 - m/z 1000) with staggered 4-m/z windows (30k resolution for precursor and product ion scans, all in the orbitrap). The 4-mz data files were used to create a DIA chromatogram library by searching against a Prosit-generated predicted spectral library based on the UniProt_Homo sapiens reviewed library_20191022.^21, 22^ Experimental samples were blocked by subject and randomized within each block. Injections of 2 µg of peptides were employed. MS data for experimental samples were acquired in the orbitrap using 8-m/z windows (staggered; 30k resolution for precursor and product ion scans) and searched against the chromatogram library. Scaffold DIA (v2.1.0; Proteome Software) was used for all DIA data processing. Results were filtered at 1% FDR (protein). DSP and desmosomal proteins were normalized to total myosin peptide counts for heart tissue.^23^

### Transcript Quantification

Human heart tissue was prepared for RNA-seq by rotor homogenization in Qiagen RLT buffer and RNA was isolated using the RNAeasy kit (Qiagen). Each biologic replicate of hPSC-CMs for RNA-seq was obtained from a separate batch of cardiac differentiation. All samples used for RNA library preparation had an RNA abundance >25 ng/µl, 260/280 ratio >1.9 (by NanoDrop^TM^), and RNA integrity number >8.0. A stranded and barcoded RNA library was prepared and sequenced using a multiplexed library on an Illumina NovaSeq by the University of Michigan DNA Sequencing Core. Reads were aligned to the genome using STAR.^24^ Lowly expressed genes were filtered. Reads per gene were normalized using DESeq2.^25^ Relative abundance for each gene was determined using dispersion estimates for each gene fit to a binomial generalized linear model with DESeq2.^25^ *DSP* transcript variant proportions were calculated using the number of unique exon 23-24 splice junctions for each transcript variant, and Sashimi plots were generated using the Integrative Genomics Viewer.^26^ Transcriptomics data were analyzed and visualized using Advaita Bioinformatic’s iPathwayGuide.^27^ For gene set analyses, a p-value <0.05 and log-2 fold change >0.2 were used to designate genes as differentially expressed. These data were deposited in NCBI’s Gene Expression Omnibus (GEO accession in progress). Quantification of mRNA for *DSP* for CRISPRi and CRISPRa experiments was performed using a TaqMan^TM^ assay from ThermoFisher (Hs00950591_m1).

### CRISPR-Cas9 Transcriptional Inhibition and Activation

CRISPick (Broad Institute) was used to select gRNA target sites (**Supplemental Table**).^28^ Desmoplakin activation/repression was performed at differentiation day 17 while cells were plated in 24 well plates. The dCas9 lentiSam V2 system was used for *DSP* activation (Addgene plasmids 75112, 89308).^29^ CRISPRi was also performed using lentiviral expression constructs (Addgene 99372, 99377). Lentivirus was prepared by the University of Michigan Vector Core.

For activation, equal amounts of dCas9 and helper or helper only lentivirus were transduced into iPSC-CMs; for repression equal amounts of dCas9-KRAB and guide RNA lentivirus were transduced into iPSC-CMs. Cells were selected with hygromycin only (control) or hygromycin + blasticidin (activation/repression) for four days. Prior to replating to CMBs, cells were cultured without antibiotics for an additional 24 hours. For qRT-PCR assays, RNA was collected after 7 days.

### Cardiac Muscle Bundles and Cardiomyocyte Adhesion Assay

Cardiac muscle bundle substrates were fabricated and hPSC-CMs replated onto these substrates as previously described.^18^ A 25:75 RPMI:DMEM mixture with 1X B27 supplement was used to raise calcium to 1.45 mM, as previously described.^18^ CMBs were assessed after 8 additional days (i.e. day 32 post-differentiation). CMBs were assayed in their regular media at 37°C and 5% CO_2_ for live cell experiments. Microscopy was performed using a Nikon Eclipse Ti-E inverted microscope. CMBs were selected and time-series images were obtained at 30 frames per second as described.^18^ Contractile function was quantified from brightfield images using our prior method, ContractQuant.^18^

For the cardiomyocyte cell-cell adhesion assay, 10 nM endothelin-1 (ET-1, Sigma, E7764) was added following baseline imaging. ET-1 was prepared in single use aliquots and stored under nitrogen gas at −20°C. CMBs were re-imaged every hour for up to 4 hours after ET-1 treatment. In a subset of experiments mavacamten (500 nM, Sigma) was used to inhibit contractility and added after baseline imaging and prior to ET-1 addition. After time series images were obtained baseline to post ET-1 time points were compared in a blinded fashion. Adhesion failure was defined as presence of cell-cell breaks or cell loss in the ET-1 treated CMBs compared to baseline.

### Immunofluorescence Imaging and Desmosome Quantification

CMBs were fixed with 4% paraformaldehyde. Desmosomes were either visualized directly (DSP-GFP reporter line and its DSP model derivatives, *DSP*^tr1-/+^ and *DSP*^tr1-/tr1-^) or by first immunostaining with a DSP antibody (1:400, Protein Tech 68364-1-Ig) followed by a goat anti-mouse AlexaFluor 488 secondary (1:1000, ThermoFisher A-11001). Adherens junctions were labeled using an N-cadherin antibody (1:400, BD Biosciences 610921) and goat anti-mouse AlexaFluor 647 (1:1000, ThermoFisher A-21235) or 568 secondary (1:1000, ThermoFisher A11004). Myofibrils were labeled using either an antibody to alpha-actinin (1:1000, Sigma A7811) followed by goat anti-rabbit AlexaFluor 568 secondary (1:1000, ThermoFisher A11036), or by direct labeling using the F-actin cell permeant dye, SiR-Actin (0.5 µM for 1 hour, Cytoskeleton, Inc.).

Fluorescence images were obtained as a z-stack with 1-µm slice thickness, and deconvolutions with maximum intensity projections were performed in Nikon Elements. Quantification of DSP intensity was performed by first creating a binary mask at the locations of N-cadherin signal; the sum intensities of desmoplakin co-localized to N-Cadherin signal were then acquired.

## Results

### Human Hearts with DSPtv Exhibit Haploinsufficiency, Desmosomal Disruption, and Excessive Fibrosis Compared to both Control and TTNtv Hearts

The majority of patients with DSP cardiomyopathy have a *DSP*tv, but studies that definitively establish haploinsufficiency in human heart tissue have been limited. Therefore, we obtained heart tissue at the time of heart transplant from patients with either *DSP*tv or *TTN*tv cardiomyopathy presenting with similar extents of left ventricular remodeling, as well as from donor heart controls (**Table 1**). *DSP*tv tissues were from females, reflecting the female predominance of this condition.^1^ In RNA-seq analysis, *DSP*tv hearts exhibited a reduction in *DSP* mRNA, consistent with nonsense mediated RNA decay (**Figure 1A**). Gene ontology analysis of transcriptomic data for *DSP*tv vs *TTN*tv revealed a significant over-representation of genes in biologic pathways associated with inflammatory response, leukocyte proliferation, cytokine production, ECM-receptor interactions, macrophage activation, and cell adhesion (**Figure 1B**). Notably, genes driving the significant differences in *DSP*tv hearts for inflammation and immune pathways were generally also upregulated in *TTN*tv compared to control hearts, but to a lesser extent (**Figure 1C**).

**Figure 1.**
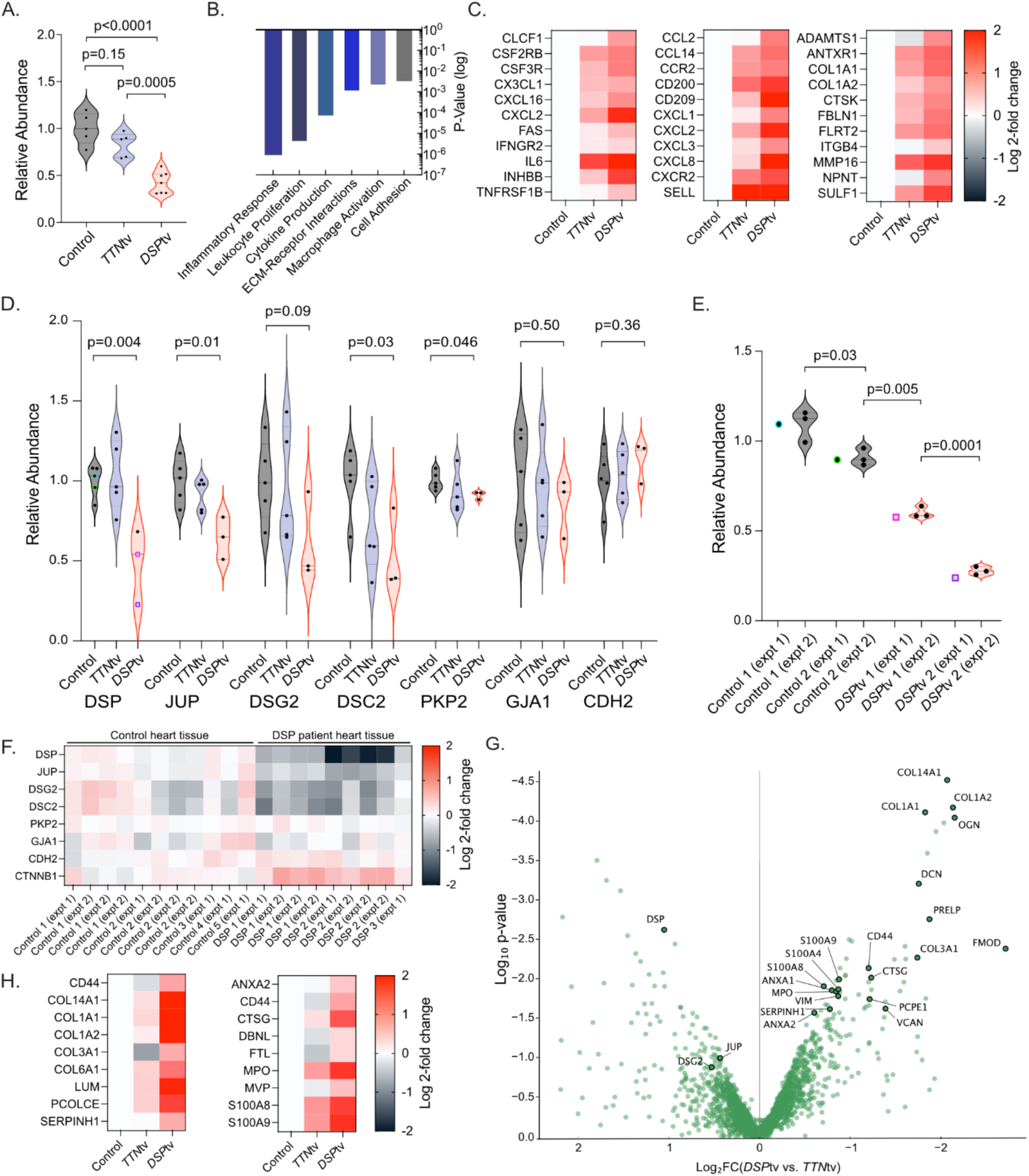
Heart tissue from patients with *DSP* truncating variants exhibit DSP haploinsufficiency and desmosomal stoichmetric disruption. A. *DSP* mRNA was quantified by RNA-seq *DSP*tv patients (N=7 separate samples from 3 patients), *TTN*tv patients (N=5), and donor heart controls (N=5). **B.** Transcriptomic gene ontology analysis was performed for *DSP*tv vs *TTN*tv patient explanted heart tissue to distinguish effects of *DSP* truncating variants in similarly remodeled left ventricular tissue. **C.** Representative genes differentially expressed in *DSP*tv vs *TTN*tv heart tissue driving differences in the cytokine (left), macrophage (center), and extracellular matrix (right) gene sets are shown. *TTN*tv heart tissue exhibited up-regulation of many of these genes compared to donor heart controls, but with lesser magnitude than *DSP*tv tissue. **D.** Mass spectrometry was performed in *DSP*tv (N=3), *TTN*tv (N=5), and control heart (N=5) tissue samples. Quantification of DSP, desmosomal proteins, connexin-43 (GJA1), and N-cadherin (CDH2) is shown. **E.** A second mass spectrometry experiment was performed to analyze inter-sample and regional variability for *DSP*tv (N=2 individuals with 3 additional samples) and control (N=2 individuals with 3 additional samples). The additional samples were from regionally remote tissue (septal, anterior, and lateral LV). DSP abundance is shown from the first experiment (brightly colored point, matching the same point shown in panel D) compared to the second experiment (black points) for each individual. **F.** Desmosomal, adherens junction, and gap junction protein quantifications are shown across all samples from both mass spectrometry experiments by heat map. **G.** Proteomic analysis was performed for *DSP*tv vs *TTN*tv heart tissue and shown by volcano plot. **H.** Proteomics ontology analysis revealed up-regulation of extracellular matrix (left) and neutrophil degranulation (right) gene sets; proteins from these biologic function categories are also highlighted in the volcano plot (G).

**Table 1.** Patient characteristics for heart tissue analyses. *This patient carried an additional *DSP* VUS (c.943C>T, p.Arg315Cys). ^This patient carried an additional *DSP* VUS (c.7885G>A, p.Glu2629Lys) and a pathogenic variant in *TNNT2* (c.629_631delAGA, p.Lys210del).

| Gene | Designation | cDNA Variant | Protein Variant | Age | Sex | LVEF |
| --- | --- | --- | --- | --- | --- | --- |
| **Control** | Control 1 | NA | NA | 18 | F | 70 |
| **Control** | Control 2 | NA | NA | 57 | F | 70 |
| **Control** | Control 3 | NA | NA | 51 | F | 65 |
| **Control** | Control 4 | NA | NA | 59 | M | 60 |
| **Control** | Control 5 | NA | NA | 60 | M | 60 |
| ***TTN*** | *TTN*tv 1 | c.43360C>T | p.Arg14454* | 69 | M | 10 |
| ***TTN*** | *TTN*tv 2 | c.51436+1G>A | NA | 39 | M | 10 |
| ***TTN*** | *TTN*tv 3 | c.73750C>T | p.Gln24584* | 44 | M | 10 |
| ***TTN*** | *TTN*tv 4 | c.53393delG | p.Gly17798fs | 45 | F | 10 |
| ***TTN*** | *TTN*tv 5 | c.35314G>T | p.Glu11772* | 54 | M | 25 |
| ***DSP*** | *DSP*tv 1 | c.491_492delins15 | p.Ala164fs | 60 | F | 15 |
| ***DSP*\*** | *DSP*tv 2 | c.1146delT | p.Phe382fs | 21 | F | 15 |
| ***DSP*<sup>^</sup>** | *DSP*tv 3 | c.7491_7492delTG | p.Cys2497* | 62 | F | 10 |

We next assessed DSP and associated desmosomal protein levels by mass spectrometry. *DSP*tv heart tissue exhibited a reduction in DSP protein levels compared to both control and *TTN*tv heart tissue (**Figure 1D**). Other desmosomal proteins, including plakoglobin (JUP), desmocollin-2 (DSC2), and plakophilin-2 (PKP2) were also significantly reduced in *DSP*tv heart tissue (**Figure 1D**). The adherens junction protein N-cadherin (CDH2) was not different in *DSP*tv heart tissue (**Figure 1D**). Noting significant variability in DSP levels among the *DSP*tv samples, we next sought to determine whether this variability is due to biologic variability, tissue sampling variability, and/or technical variability. Therefore, we repeated the mass spectrometry experiment using multiple separate samples from hearts for which regional tissue was available (i.e., septal, anterior, and lateral left ventricle samples from N=2 *DSP*tv, N=2 control). We observed minimal differences in DSP levels from the initial experiment and across regional samples measured in the second experiment (**Figure 1E**). Interestingly, an 18% difference in DSP level was present across the two control hearts (p=0.03, **Figure 1E**). DSP levels were markedly reduced in both *DSP*tv hearts with a significantly larger reduction in the tissue from patient *DSP*tv 2 (39.8±3.1% and 72.2±2.3%, p=0.0006 and p<0.0001 vs. controls; p=0.0001 for *DSP*tv 1 vs. *DSP*tv 2, **Figure 1E**). Notably, the *DSP*tv 2 heart tissue was collected from a patient with much earlier progression to advanced heart failure (heart transplant at age 21 vs age 60); this patient also carried a missense variant in *DSP* on the other allele (**Table 1**). We observed a similar pattern of reductions in desmosomal proteins across both mass spectrometry experiments (**Figure 1F**). From experiment 1, we performed a proteomic analysis to compare *DSP*tv to *TTN*tv heart tissue (**Figure 1G-H**). Pathway analysis revealed significant enrichment of dysregulated proteins in integrin cell surface interactions (p=5.5e-18), collagen biosynthesis (5.3e-17), and neutrophil degranulation (1.2e-16). Genes driving these differences are highlighted in the volcano plot (**Figure 1G**) and heat maps (**Figure 1H**). Several proteins in these pathways exhibited similar directionality (but lower magnitude) of effect in *TTN*tv heart tissue when compared to control heart tissue (**Figure 1H**). Notable proteins that were uniquely up-regulated in *DSP*tv heart tissue were ANXA1 (p=0.004), ANXA2 (p=0.03), and CD44 (p=0.03).

### iPSC-CM Models with DSPtv Recapitulate Haploinsufficiency, Desmosomal Disruption, and Dysregulation of Cell Adhesion Genes

We next developed experimental models of DSP cardiomyopathy using iPSC-CMs. Our goal was to model severe DSP haploinsufficiency such that iPSC-CMs would capture relevant disease pathology over the time course feasible for cell culture, but without resorting to complete DSP knock-out, which may reflect pathologies not relevant for the human disease. Therefore, we generated patient derived iPSC lines from one of the patients who underwent heart transplant (“*DSP*tv 2” in **Table 1**), in addition to two of this patient’s siblings (one also severely affected and one mildly affected, **Figure 2A**). Additionally, we gene edited a control DSP-GFP reporter iPSC line to have both single and biallelic truncating variants exclusively in *DSP* transcript 1, which encodes the dominant cardiac DSP isoform, DSPI (**Figure 2B**). Based on RNA-seq quantification of control human heart tissue (N=5, as above), *DSP* transcript 2 (encoding DSPII) accounts for 22.7±2.3% of total cardiac *DSP* expression (**Supplemental Figure 2**). Importantly, patients born with biallelic DSPI truncating variants live into childhood before developing severe heart failure, as opposed to total DSPI/II loss of function, which results in embryonic lethality^30, 31^ These lines, designated as *DSP*^tr1-/+^ and *DSP*^tr1-/tr1-^, were expected to exhibit levels of *DSP* mRNA expression of 62% and 23% of normal, respectively (**Figure 2B**).

**Figure 2.**
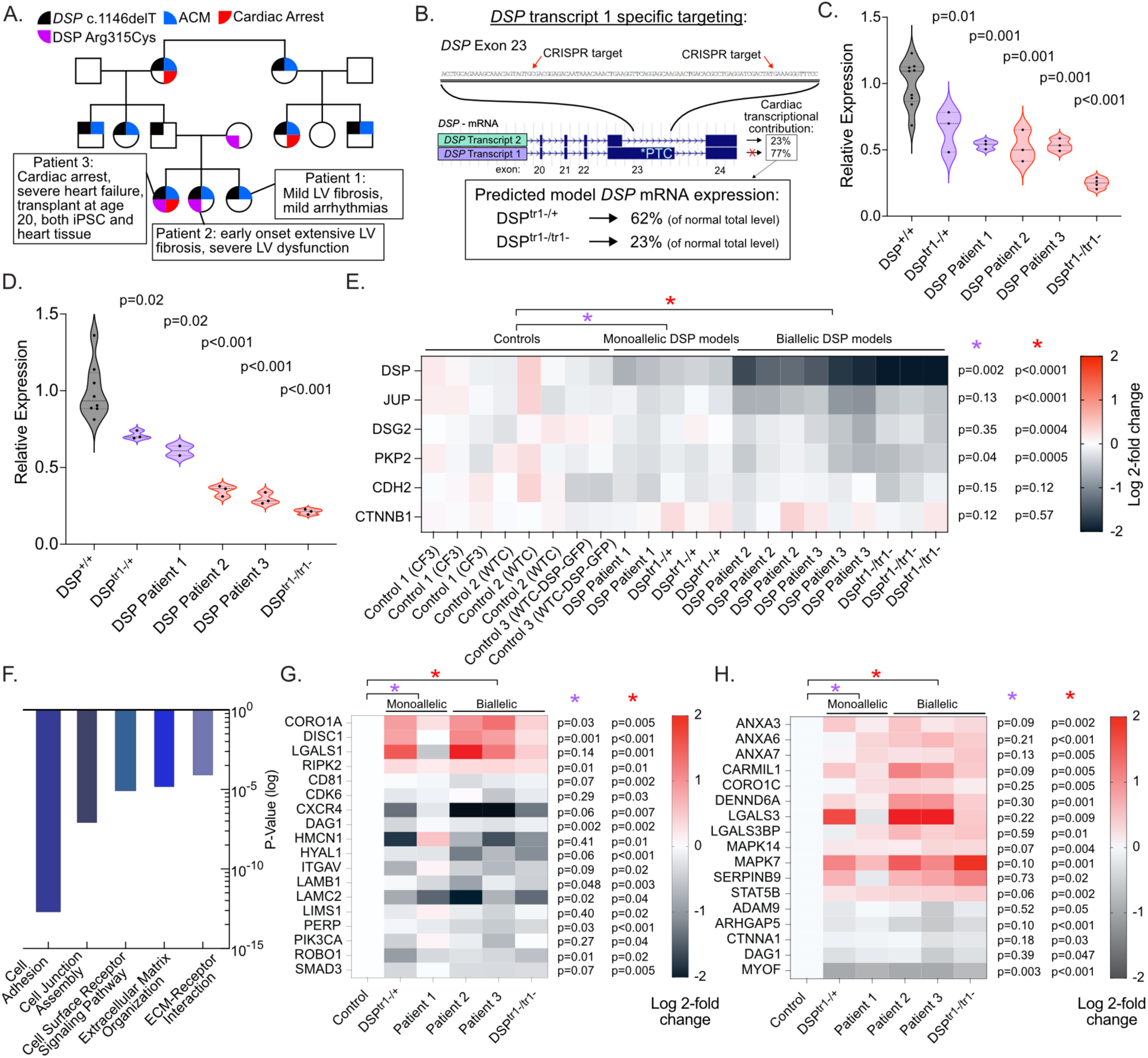
*DSP*tv iPSC-CM models recapitulate DSP haploinsufficiency, desmosomal stoichiometric disruption, and dysregulated cell adhesion genes. A. Family pedigree is shown for *DSP*tv patients for whom iPSC-CMs were generated. Patient 3 is the same individual as for whom explanted heart tissue was collected (DSP 2 in Figure 1E-F). *DSP* c.1146delT is the pathogenic *DSP*tv. DSP Arg315Cys was designated as a variant of unknown significance and was also present in both severely affected patients in tandem with the *DSP*tv. B. *DSP* transcript 1 was targeted by introducing frameshift variants in the portion of exon 23 that is only spliced into *DSP* transcript 1. Monoallelic (*DSP*^tr1-/+^) or biallelic (*DSP*^tr1-/tr1-^) targeting were predicted to result in residual *DSP* mRNA expression of 62% and 23% respectively (being exclusively from *DSP* transcript 2), based on our splicing quantification from human heart tissue (Supplemental Figure 2). These iPSC lines were made on the WTC-DSP-GFP reporter line background. C. *DSP* mRNA was quantified by RNA-seq from 3 separate differentiation batches for each iPSC-CM line. **D.** DSP quantification was performed by mass spectrometry from 3 separate differentiation batches for each iPSC-CM line. **E.** Desmosomal and adherens junction protein quantifications are shown across all samples from both mass spectrometry experiments by heat map. Statistical tests (ANOVA) are shown comparing iPSC-CM lines with severe DSP reduction due to biallelic variants (*DSP*^tr1-/tr1-^, Patient 2, Patient 3) or moderate reduction (*DSP*^tr^^1-/+^, Patient 1) to control lines. **F.** Transcriptomic gene ontology analysis was performed for *DSP*tv vs. control iPSC-CMs with representative statistically significant gene categories shown. **G.** Cell adhesion genes differentially expressed in *DSP*tv iPSC-CMs across all models are shown by heat map (see Supplemental Figure 3 for network diagram of these and related genes). **H.** Proteomics ontology analysis revealed differential protein levels for cell adhesion and membrane function related proteins; several proteins driving these differences are shown by heat map.

We characterized this panel of DSP-CM models initially with RNA-seq and mass spectrometry using the same methodology as for human heart tissue. Similar to *DSP*tv human hearts, we found that *DSP* mRNA was reduced across all models (**Figure 2C**). As expected, *DSP*^tr1-/tr1-^ iPSC-CMs retained 25±4% residual expression due to intact *DSP* transcript 2 (**Figure 2C**).

Mass spectrometry quantification demonstrated a reduction in DSP protein across all models, again with severe reduction in *DSP*^tr1-/tr1-^ iPSC-CMs (residual 21±2% of normal, p<0.0001, **Figure 2D**). The two iPSC-CM lines derived from severely affected patients (Patient 2 and Patient 3, Figure 2A) also exhibited more severe reductions in DSP protein (residual 35±3% and 30±4% of normal, respectively) – the latter of these approximating the reduction in DSP measured from heart tissue for the same patient (see above). Notably, these two patients both carried a *DSP* variant of unknown significant (VUS) on their other *DSP* allele, putatively acting as a hypomorphic allele. Hence, these iPSC-CM lines were considered as biallelic models, along with the *DSP*^tr1-/tr1-^ line. Mass spectrometry analysis of other desmosomal proteins revealed a similar pattern of reduced levels as had been observed in human heart (**Figure 2E**). Specifically, plakoglobin (JUP), desmoglein-2 (DSG2), and plakophilin-2 (PKP2) were significantly reduced in the biallelic DSP iPSC-CM models. Also, as in human *DSP*tv heart tissue, the adherens junction protein N-cadherin (CDH2) was unaltered. Of note, GJA1 and DSC2 levels by mass spectrometry were more variable in control iPSC-CMs and therefore are not shown in Figure 2E due to lower confidence (standard deviation 0.76 and 0.98, respectively; all proteins shown in Figure 2E had standard deviations ≤0.2).

Transcriptomic gene ontology analysis revealed over-representation of pathways related to cell adhesion, cell junction assembly, and extracellular matrix organization (**Figure 2F**). Dysregulated genes driving these pathways are shown in **Figure 2G**. Adhesion-related genes included *CXCR4*, *CXCL12*, *ROBO1* (which regulates CXCR4-CXCL12 interactions), PERP, laminin genes (e.g. *LAMB1*, *LAMC2*), and integrin genes (e.g. *ITGAV, LIMS1*). A network diagram of differentially expressed cell adhesion genes is shown in **Supplemental Figure 3**. Proteomic analysis of iPSC-CM models revealed that dysregulated proteins primarily localized to extracellular space (p=0.0007) or plasma membrane (p=0.002). Over-represented pathways included cell communication (p=0.0009), cell migration (p=0.0009), desmosome organization (p=0.002), and cell adhesion molecule (p=0.004). Selected proteins from these pathways are shown in **Figure 2H**. Notable up-regulated proteins included membrane injury-repair associated annexins (ANXA3, ANXA6, ANXA6) and the injury response mediators, galectin-3 (LGALS3) and p38⍺ (MAPK14). Notable down-regulated proteins included dystroglycan (DAG1) and myoferlin (MYOF), both involved in normal cardiomyocyte membrane function.

Taken together, these analyses demonstrate that *DSP* truncating variants acutely result in DSP protein level haploinsufficiency, altered desmosomal stoichiometry, and broad disruption of genes/proteins involved in cell-cell adhesion, membrane function, and cardiomyocyte response to injury.

### DSPtv Cardiac Muscle Bundles Retain Normal Baseline Structure and Contractile Function but Exhibit Cell-Cell Adhesion Failure with Heightened Contractile Stress

To assess the impact of DSP truncating variants on cardiomyocyte function, we used a previously established platform for generating 2D cardiac muscle bundles (CMB).^18^ We selected the CMB platform since it enables greater uniaxial contractile force generation in micropatterned iPSC-CM tissues on an elastomeric substrate that replicates cardiac tissue stiffness (∼8 kPa).^18^ This approach results in robust alignment of myofibrils with intercalated disk protein complexes localized at cell-cell junctions. Moreover, CMBs are a single cell layer thick and therefore conducive to live cell visualization of cell-cell junctions (as we previously demonstrated with both DSP-GFP and connexin-43-GFP reporter iPSC-CMs).^18^ To increase the signal to noise ratio for functional analyses, we focused these studies on the two most severely haploinsufficient *DSP*tv models (*DSP*^tr1-/tr1-^ and *DSP* Patient 3).

*DSP*tv iPSC-CMs assembled normally into CMBs (**Figure 3A and B; Supplemental Figure 4**). Immunofluorescence imaging demonstrated myofibrillar alignment (marked by ⍺-actinin) along the tissues’ long axes with comparable organization between control and *DSP*tv models. DSP localized to cell junctions and co-localized with the adherens junction protein, N-cadherin, similarly in all models (including *DSP*^tr1-/tr1-^ iPSC-CMs, which express only *DSP* transcript 2), although with an evident diminished abundance in *DSP*tv models (**Figure 3A and B; Supplemental Figure 4**). CMBs exhibited regularity of organized spontaneous contractions for all controls/models (**Figure 3C-D**). Contractile function was variable between control lines, as we have previously observed in both single cell iPSC-CM and CMB contractile studies (**Figure 3E-F**).^18, 32^ Neither contractile velocity nor maximal fractional shortening were reduced in *DSP*^tr1-/tr1-^ CMBs compared to isogenic control (WTC-*DSP*-GFP) CMBs (**Figure 3E**-**F**). Similarly, DSP Patient 3 CMBs did not exhibit reduced contractile velocity or reduced maximal fractional shortening compared to the additional control line (**Figure 3E-F**).

**Figure 3.**
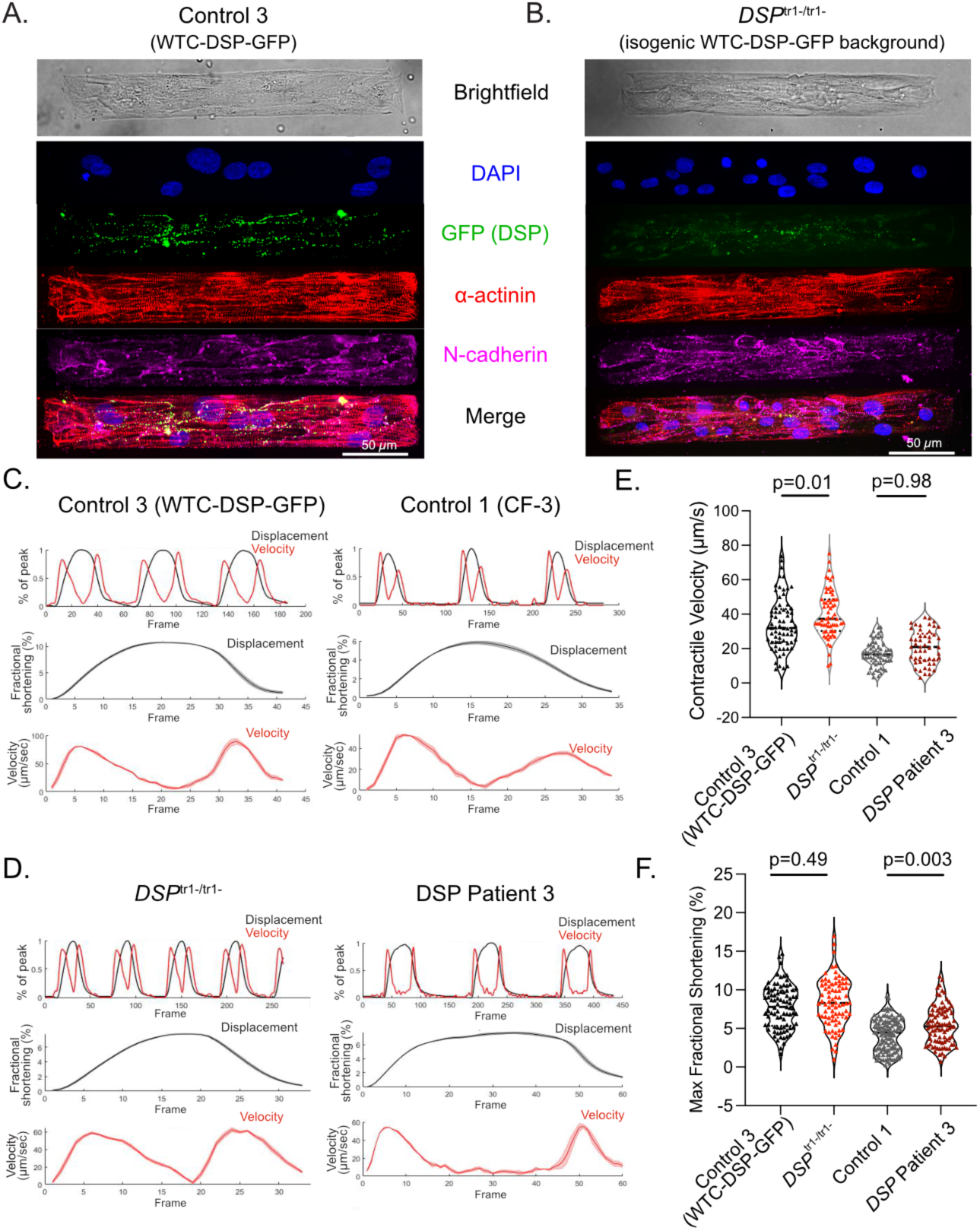
*DSP* haploinsufficient cardiac muscle bundles (CMBs) do not exhibit overt structural or functional defects under baseline conditions. A-B. Brightfield and immunofluorescent images of CMBs generated from control 3 (WTC-DSP-GFP) and *DSP*^tr1-/tr1-^ (isogenic WTC-DSP-GFP background) iPSC-CMs. Immunofluorescent images showing DAPI (blue, nuclei), GFP (green, DSP), α-actinin (red, sarcomeres), and N-cadherin (magenta, intercalated discs/cell junctions), demonstrating comparable morphology and protein localization at baseline. **C.** and **E.** Representative output from ContractQuant for CMBs showing (top) concurrent normalized displacements and velocities for the full time-series capture (second row) fractional shortening of merged contractions, (third row) merged absolute values of contractile velocity of control and *DSP*tv cardiomyocytes. **D.** Contractile velocities of Control 3, *DSP*^tr1-/tr1-^, Control 1, and *DSP* Patient 3 CMBs in baseline conditions, two-sided P value shown from unpaired t test. **F.** Max fractional shortening of Control 3, *DSP*^tr1-/tr1-,^ Control 1, and DSP Patient 3 CMBs in baseline conditions, two-sided P values shown from unpaired t test, data are presented as violin plots showing individual data points, median, and interquartile ranges. All contractile experiments were pooled from at least three distinct experiments, CMBs for each experiment were generated from a unique differentiation batch.

We next hypothesized that reduced DSP expression in the *DSP*tv models would render them susceptible to cell adhesion failure under heightened contractile stress. To test this hypothesis, we established conditions of heightened contractile stress. We found that endothelin-1 (ET-1), a G_q_ protein activator, is a reproducible and potent contractile agonist in CMBs and increased contractile frequency, maximum fractional shortening, and contractile velocity acutely in both control and *DSP*tv models (**Figure 4A-C**).^33^ However, upon prolonged exposure to ET-1 (i.e. 2 hr), *DSP*tv CMBs exhibited cell-cell adhesion failure whereas control CMBs remained intact (**Figure 4D, 4F, Supplemental Figure 5, Supplemental Videos 1-3**). Next, to test whether adhesion failure could be prevented by blunting the hypercontractile response of ET-1, we pre-treated *DSP*tv CMBs with the cardiac myosin inhibitor, mavacamten. Mavacamten-treated *DSP*tv CMBs did not exhibit adhesion failure (**Figure 4E, 4F**).

**Figure 4.**
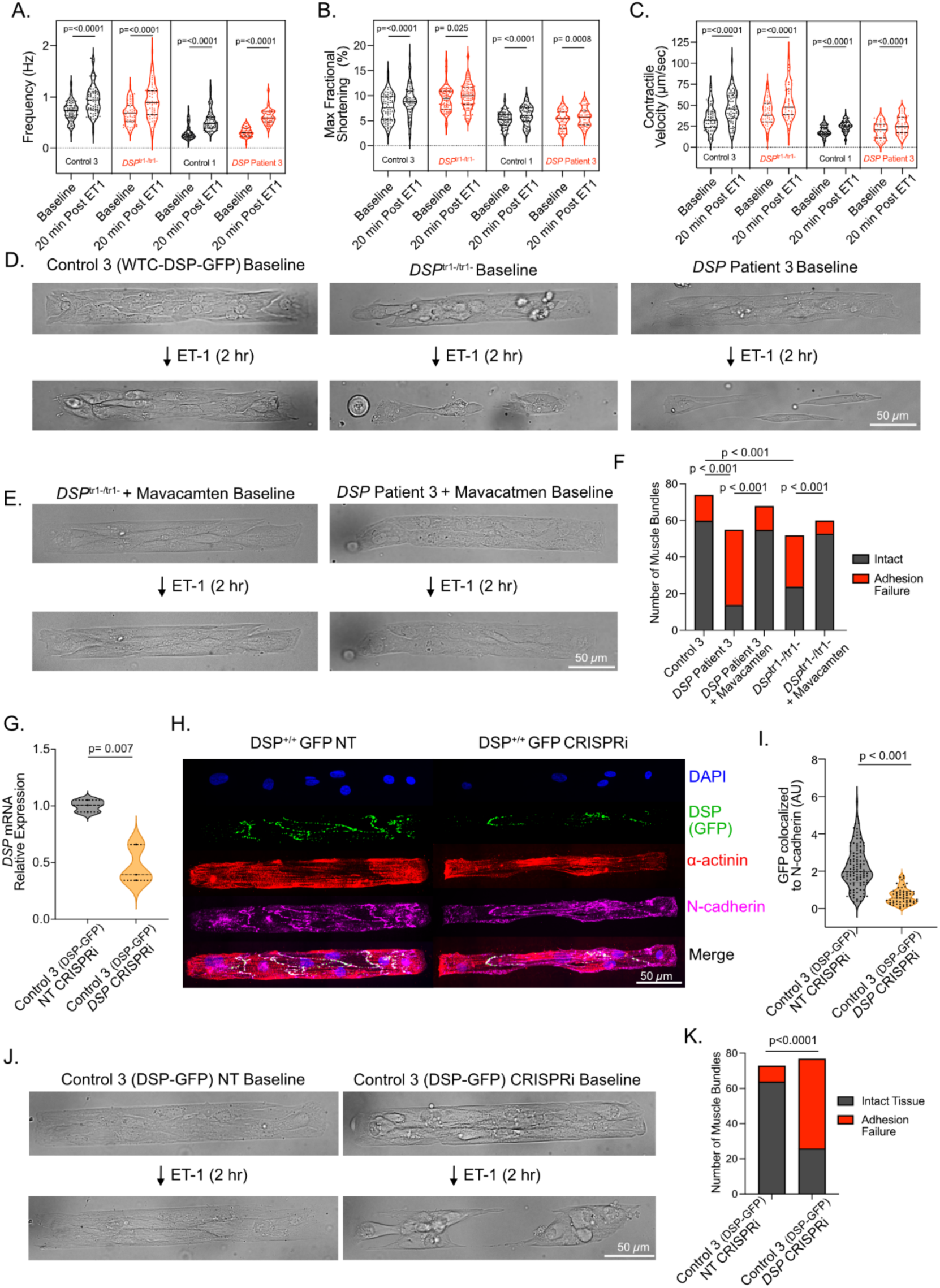
An endothelin-1 (ET-1) cardiac muscle bundle (CMB) stress assay reveals cell-adhesion failure in DSP-deficient (DSPtv) cardiomyocytes and in control cardiomyocytes after CRISPRi-mediated DSP transcriptional repression. **A-C**. ET-1 (10 nM) elicits a positive inotropic and chronotropic response in CMBs from all tested lines: Control 3, *DSP*^tr1-/tr1^, Control 1, *DSP* Patient 3, demonstrating increased contractile frequency (A), maximum fractional shortening (B), and contractile velocity (C) 20 minutes post-treatment, thereby validating ET-1 as an effective stressor imposing heightened mechanical load. Two-sided P values shown from paired t tests. **D**. Representative brightfield images of CMBs from Control 3, *DSP*^tr1-/tr1-,^ and *DSP* Patient 3 lines at baseline and after 2 hours of continuous ET-1 (10nM) exposure. CMBs from DSPtv lines exhibit pronounced cell-cell adhesion failure under sustained ET-1 induced stress. **E.** Treatment with Mavacamten (500 nM) prior to ET-1 addition prevents ET-1 induced cell-adhesion failure in *DSP*^tr1-/tr1-^ and *DSP* Patient 3 CMBs. Representative brightfield images are shown at baseline (with Mavacamten) and after 2 hours of ET-1 exposure in the continued presence of Mavacamten. **F**. Quantification of CMB integrity after 2-hours of ET-1 stress, with or without Mavacamten treatment. The bar graph shows the number of intact CMBs versus those exhibiting adhesion failure for the various cell lines tested. **G**. Quantitative PCR analysis shows significantly reduced *DSP* mRNA levels in CMBs transduced with *DSP*-targeting CRISPR-Cas9 interference (CRISPRi) vs. non-targeting CRISPRi controls. **H.** Representative immunofluorescence images demonstrating reduced GFP-tagged DSP protein (green) at cell junctions (N-cadherin, magenta) in *DSP* CRISPRi CMBs versus NT CRISPRi CMBs. **I.** Quantification of GFP (DSP) signal intensity colocalized with N-cadherin at cell-cell junctions, confirming significant DSP protein knockdown in *DSP* CRISPRi CMBs compared to NT CRISPRi controls. **J**. Representative brightfield images of Control 3 (WTC-DSP-GFP) CMBs transduced with NT CRISPRi or *DSP* CRISPRi, shown at baseline and after 2 hours of ET-1 (10nm) exposure. *DSP* CRISPRi CMBs exhibit adhesion failure upon ET-1 stress, while NT CRISPRi CMBs remain intact. **K**. Quantification of CMB integrity after 2-hour ET-1 stress in CRISPRi-treated control cells. The bar graph shows a significantly higher percentage of adhesion failure in *DSP* CRISPRi CMBs compared to NT CRISPRi CMBs when exposed to ET-1. All data were generated from at least 3 independent experiments and cardiomyocyte differentiations. Data are presented as violin plots showing individual data points, median, and interquartile ranges. Continuous variables were compared with two-sided unpaired t-tests, except for **A, B,** and **C.** which used two-sided paired t-tests. Categorical variables were compared with Fischer’s Exact Test.

Collectively, these results demonstrate that *DSP*tv iPSC-CMs assemble normally into organized muscle bundle tissues and have normal baseline contractile function. However, *DSP*tv CMBs demonstrate marked sensitivity to cell-cell junction failure when challenged with heightened contractile stress.

### Transcriptional Repression with CRISPRi Results in Cell-Cell Adhesion Failure Similarly as in DSPtv models

To mechanistically determine if DSP haploinsufficiency causes cardiomyocyte cell-cell adhesion failure, we next generated CRISPRi constructs. CRISPRi targeted to the *DSP* promoter reduced *DSP* mRNA of control cells by 53±10% (p=0.006, **Figure 4G**) and DSP protein levels by 69±9% (p<0.0001, **Figure 4H-I**). *DSP* CRISPRi CMBs assembled normally and demonstrated similar baseline contractility to control (non-targeted CRISPRi) CMBs (**Figure 4J, Supplemental Figure 5 and 6**). However, after exposure to ET-1, *DSP* CRISPRi CMBs demonstrated adhesion failure similarly as in *DSP*tv CMBs (**Figure 4J-K**, **Supplemental Figure 5**). These experiments demonstrate that *DSP* loss of function is sufficient to cause cell-cell adhesion failure with contractile stress.

### Transcriptional Activation with CRISPRa Rescues DSP Levels and Cell-Cell Adhesion

Having demonstrated DSP haploinsufficiency in human heart tissue and mechanistically proven that DSP loss of function drives a primary defect of cell-cell adhesion failure, we next tested whether this pathology could be rescued by transcriptional activation of *DSP*. Targeting three sites in the *DSP* promoter with different guide RNAs showed each site could effectively drive transcriptional activation of the *DSP* gene (**Supplemental Figure 7**). We further validated *DSP* transcriptional up-regulation independently in two control and the two most severely haploinsufficient *DSP*tv iPSC-CM lines (**Figure 5A**). Both wild type and *DSP*tv iPSC-CMs demonstrated a similar relative magnitude of increase (**Figure 5A**, controls: *DSP^+/+^* GFP 83% increase, p<0.001, *DSP*^+/+^86% increase, p=0.002, *DSP*^tr1-/tr1-^ 82% increase, p<0.001, and DSP patient 3 86% increase, p=0.004). To assess both DSP protein quantity and localization, CRISPRa CMBs were generated. As shown in **Figure 5B**, DSP successfully localized to cell junctions. Notably, our CRISPRa strategy targets the DSP promoter region and is not intended to be allele-specific. Additionally, in the *DSP*^tr1-/tr1-^ iPSC-CM model, which has biallelic loss-of-function variants affecting only DSPI, this transcriptional upregulation effectively enhances only the expression of DSPII. We did not observe mislocalized DSP protein at the levels of expression attained in either the *DSP*^tr1-/tr1-^ or *DSP* Patient 3 CMBs (**Figure 5B**). The quantity of DSP protein colocalized to N-cadherin increased in both *DSP*^tr1-/tr1-^ and *DSP* Patient 3 (50%, p<0.001 and 60%, p<0.001, respectively).

**Figure 5.**
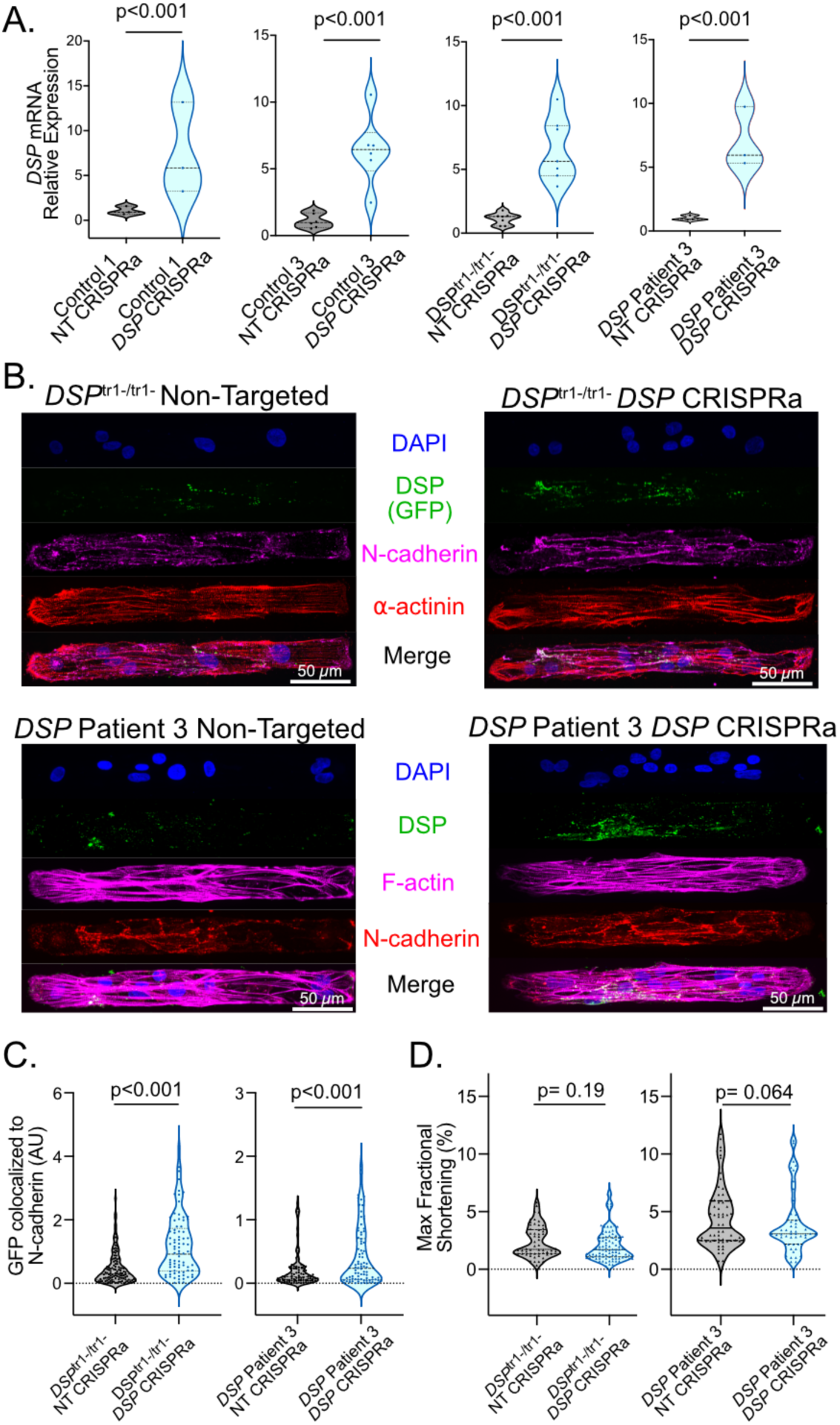
Transcriptional activation with CRISPR-dCas9 increases DSP expression and localization in control and patient-derived iPSC-CMs. **A.** Relative *DSP* mRNA expression measured by quantitative PCR in control and *DSP*tv cardiomyocytes demonstrates the ability to increase mRNA expression in both control and disease cell lines. **B**. Representative immunofluorescence images of non-targeted and CRISPRa treated *DSP*tv CMBs showing increased DSP (green) expression and correct localization to the intercalated disc as labeled by N-Cadherin (red). **C**. Quantification of DSP colocalization with N-cadherin in *DSP*tv CMBs showing increased expression of DSP in cells treated with CRISPRa. **D.** Maximum fractional shortening analysis in *DSP*tv CMBs showing no difference between non-targeted and CRISPRa CMBs. Continuous variable compared with two tailed, unpaired t tests, violin plots show median and interquartile ranges.

**Figure 6.**
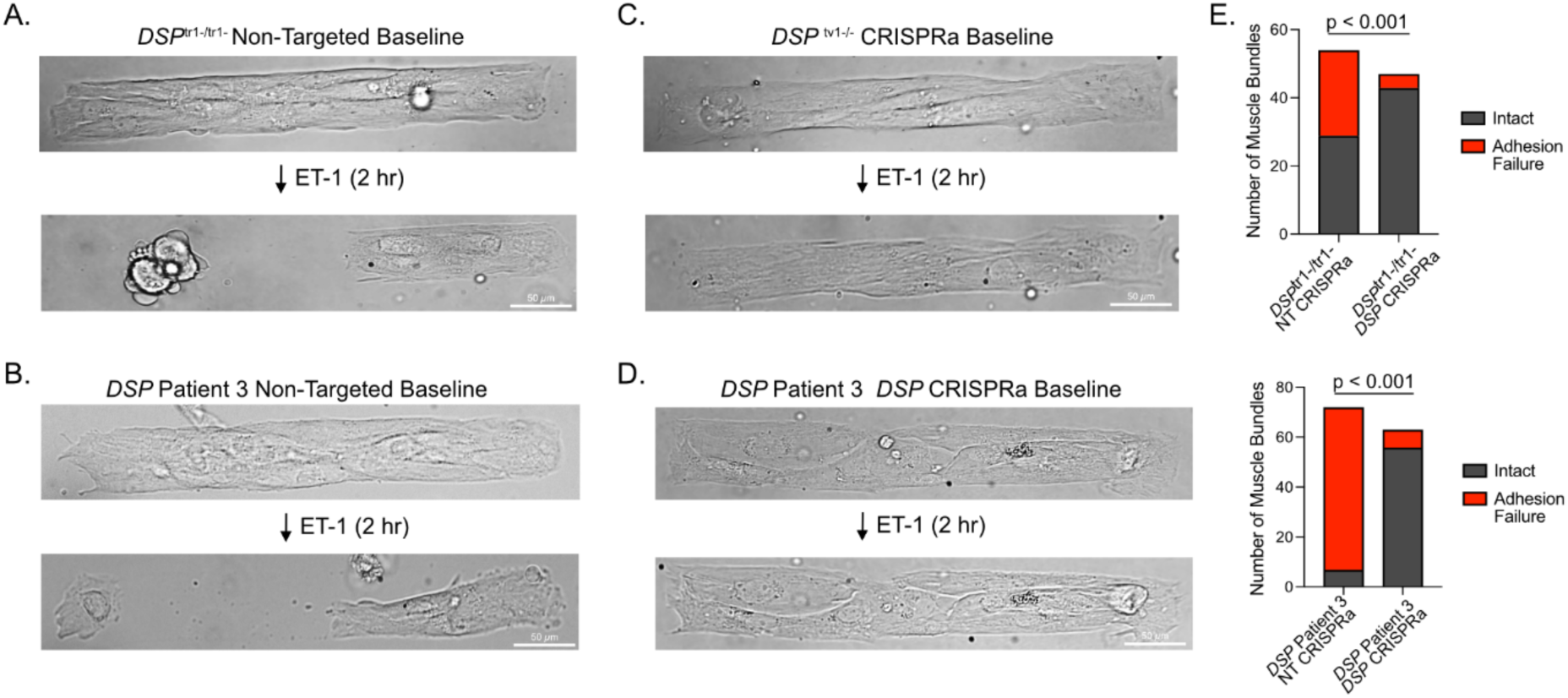
CRISPR activation of *DSP* rescues the structural integrity of cardiac muscle bundles subjected to increased contractile stress. **A-D.** Representative bright field of either non-treated (**A, B**) or CRISPRa treated (**C, D**) CMBs. Images show CMBs at baseline (top row in each panel) and after 2 hours of stimulation with endothelin-1 (ET-1, bottom row in each panel). **E.** Adhesion failure was quantified as present or absent following ET-1 challenge. Stacked bar charts show the number of bundles classified as intact (gray) or exhibiting adhesion failure (red). DSP CRISPRa treatment significantly reduced the incidence of adhesion failure in both cell lines (p < 0.001, Fisher’s exact test).

We next assessed whether increased expression of *DSP* via CRISPRa would impact CMB contractility. There was not a significant difference in contractile velocity or maximum fractional shortening between non-targeted and activated CMBs (DSP^tr1-/tr1-^ 16.3 ± 8.7 μm/s vs 14.0 ± 7.2 μm/s, p=0.07, and DSP Patient 3 22.4 ± 16.2 μm/s vs 21.1 ± 13.4 μm/s, p=0.70, **Figure 5C**), (DSP^tr1-/tr1-^ 2.4 ± 1.3% vs 2.1 ± 1.4%, p=0.18 and *DSP* Patient 3 5.7 ± 3.6% vs 4.4 ± 2.6%, p=0.06, **Figure 5D**).

Finally, we tested whether CRISPRa of DSP haploinsufficient cardiomyocytes was sufficient to rescue cell-cell adhesion failure under heightened contractile stress. *DSP*tv cardiomyocytes CMBs under control conditions (non-targeting CRISPRa) demonstrated adhesive failure under ET-1-mediated increases in contractile stress (**Figure 6A-B)**, whereas *DSP* CRISPRa resulted in marked improvements (*DSP*^tr1-/tr1-^: 46% failure vs 9%, p<0.0001; *DSP* Patient 3: 89% failure vs 11% failure; **Figure 6 C-E**). Notably, functional rescue in the *DSP*^tr1-/tr1-^ iPSC-CMs demonstrates that upregulation of DSPII is sufficient for restoration of cell-cell adhesion.

Additionally, up-regulated *DSP* transcription in Patient 3 iPSC-CMs necessarily consisted of only transcripts containing the DSP VUS Arg315Cys – these results suggest that the Arg315Cys variant may impair DSP’s expression and/or stability but not function. Taken together, these findings support the feasibility of *DSP* transcriptional activation for rescuing cell-cell adhesive dysfunction due to *DSP*tv.

## Discussion

Despite knowledge of the genetic underpinnings of ACM, immediate pathogenic consequences of causative variants have been incompletely resolved. In this study, we definitively establish DSP haploinsufficiency in explanted human *DSP*tv hearts, which also exhibited broad disruption of desmosomal stoichiometry. Additionally, using a novel platform for analyzing the function of cell-cell junctions in iPSC-CM microtissues, we found that *DSP*tv’s lead to cardiomyocyte cell-cell adhesion failure under heightened contractile stress. This mechanically induced vulnerability was mechanistically linked to DSP reduction via CRISPRi. Finally, we demonstrate that CRISPRa-mediated transcriptional upregulation is capable of restoring normal DSP protein abundance and cell adhesion function in *DSP*tv iPSC-CM models.

### Cell-Cell Adhesion Failure is a Primary Disease Mechanism for Desmoplakin Cardiomyopathy

Cell-cell adhesion dysfunction has been an intuitive theory of disease pathogenesis in arrhythmogenic cardiomyopathy due to desmosomal gene variants. This theory is suggested by altered desmosomal stoichiometry, as observed in prior murine desmosomal protein knock-out models, and as further demonstrated to be present in human desmoplakin cardiomyopathy by our mass spectrometry analyses in this study.^15, 34, 35^ Moreover, desmosomal variants, including *DSP* variants, have been linked to adhesive dysfunction in skin disease.^36, 37^ Yet, adhesive function has been challenging to directly assess in cardiac cells and tissue. One approach has been to measure relative dissociation of cardiac cells/tissue when subjected to protease digestion followed by successive pipetting, and this method has demonstrated evidence of cell-cell adhesion failure in prior desmosomal variant models.^38–40^ In particular, introduction of a *DSG2* variant (DSG2-W2A) that impairs the extracellular interaction mechanism of DSG2 molecular binding results in cell adhesive failure as determined by dissociation assays.

Specifically for DSP, studies of adhesive function have been limited. In HL-1 cells, siRNA knockdown of *DSP* did not alter cell-cell adhesion by the dissociation assay, though DSP appeared to be important for increased adhesion induced by EGFR inhibition.^41^ Bliley et al. observed an increased rate of tissue “breakages” in their EHT DSP model, implying the possibility of cell-cell adhesion loss, but the direct cellular mechanism of these events was not feasibly assessed in the EHT model.^7^

In this work, we have developed a novel method for direct visualization of cell-cell adhesion failure in iPSC-CMs utilizing a desmosome-related disease model. This method is uniquely enabled by the generation of single cell thickness cardiac muscle bundles that 1) exhibit highly aligned myofibrils in discrete cardiomyocyte tissues that contract against a controlled load imparted by the underlying elastomeric substrate, and 2) are highly amenable to high-resolution live-cell imaging.^18^ To ensure that we investigated patient-relevant disease pathology, we used iPSC-CM models that demonstrated severe (but incomplete) desmosomal loss, recapitulating findings from human heart tissue. The *DSP*^tr1-/tr-^ model, by using a DSP-GFP background line, also demonstrates that DSPII retains normal localization at cardiomyocyte junctions (with reduced abundance), explaining the survival into childhood of the rare individuals carrying biallelic *DSP*tv’s in the DSP transcript 1-specific portion of exon 23.^30, 31^ Notably, unlike dissociation assays, our method enables cell-cell adhesive function to be assessed in physiologically contracting tissues in regular media conditions. We anticipate that this assay will be of high utility for both mechanistic and translational pharmacologic studies of DSP cardiomyopathy and for extension to studies of other desmosomal diseases in the future.

### Cell Adhesion Failure May be an Upstream Driver of Cardiac Tissue Injury and Inflammation in DSP-Cardiomyopathy

Myocardial inflammation is a key contributor to desmosomal-ACM pathogenesis, and immune cell infiltration in the heart has been a consistent finding in heart tissue from patients with ACM.^10, 42^ In mouse models of DSG2-mediated ACM, activation of NF-κβ through innate immunity pathways has been shown to be a core driver of inflammation, and the capability of therapeutically targeting this pathway has also been demonstrated.^16^ Recently, NF-κβ was shown to be up-regulated in the presence of complete DSP knock-out iPSC-CM EHTs, suggesting that a similar mechanism could be involved.^8^ Our results here indicate that cell-cell adhesion failure is the likely upstream cause of cardiac injury, that then initiates an immune response and eventual reparative fibrosis. Evidence supporting this cascade was found in both our *DSP*tv iPSC-CM and heart tissue analyses. In the iPSC-CM transcriptomic and proteomic analyses, we found differentially regulated genes/proteins primarily in cell adhesion and extracellular matrix related ontologies/pathways. In particular, we observed an increase in membrane injury-repair associated annexins (ANXA3, ANXA6, ANXA6) and in galectin-3 (LGALS3), which is critical in mediating the fibrotic response to cardiac injury.^43^ Dystroglycan (DAG1), a transmembrane glycoprotein that links dystrophin to extracellular matrix and limits stress-induced membrane damage in cardiomyocytes was down-regulated in DSP models.^44^ Myoferlin, a paralog of dysferlin implicated in muscle membrane repair, was also significantly reduced in DSP models.^45^ Likewise, in *DSP*tv patient heart tissue, uniquely up-regulated proteins compared to *TTN*tv hearts included ANXA 1 and ANXA2, suggesting that muscle membrane repair pathways may remain activated in the advanced disease stage.^46^ Up-regulated proteins in *DSP*tv hearts also included CD44, the main receptor for hyaluronic acid, which is known to mediate inflammation and wound healing following myocardial injury.^47^

Another theory of desmosomal ACM pathogenesis is that the immune response, and even autoimmunity, could contribute.^48^ While our findings are more supportive of a primary role of adhesion failure leading to cardiomyocyte injury with subsequent apoptosis/necrosis and ensuing inflammatory response, these theories are not necessarily mutually exclusive. One possibility is that cell damage could result in excessive exposure to self-antigens that lead to a subsequent autoimmune response.^48, 49^ After initiation of the immune response, a feed-forward loop could ensue, promulgating further cardiac injury. In this scenario, interruption of the inflammatory response may be therapeutically advantageous beyond targeting the upstream mechanism of DSP haploinsufficiency and adhesion failure.

### Transcriptional upregulation is a potential therapeutic approach for desmoplakin cardiomyopathy

We present here definitive evidence for haploinsufficiency as the primary upstream cause of DSP cardiomyopathy due to truncating variants. Unlike the case of truncating variants in the sarcomere gene *MYBPC3*^20^, we found no evidence of significant post-translational compensation in the early developmental stage of iPSC-CMs – DSP protein levels were similarly reduced in iPSC-CMs and heart tissue collected from the same patient. The most upstream treatment for DSP cardiomyopathy would therefore be restoration of the normal level of DSP. However, the *DSP* gene is too large (8.6 kb) for the packaging limit of current AAV technology. An alternative therapeutic approach would be to upregulate *DSP* transcription. Here, we tested the concept that global transcriptional activation of *DSP* with CRISPRa can restore normal DSP levels. This approach was previously found to be efficacious for haploinsufficiency-mediated obesity caused by loss of function variants in the gene *Sim1*, and, more recently, for *TTN*tv cardiomyopathy.^50, 51^ Nonetheless, this approach does not necessarily extend to other genes, since the loss of function mechanism and the tolerated gene-dose therapeutic window requires gene-specific assessment. A potential negative consequence of this approach is that both alleles are presumably transcriptionally upregulated. Therefore, the approach assumes that the normal cellular nonsense-mediated decay process will clear all excess transcripts from the TV-containing allele. At the levels of expression we attained in our *DSP*tv models, we did not observe evidence of toxicity from overexpression – specifically, contractile function was similar with CRISPRa treatment, and we did not observe mis-localized DSP protein following CRISPRa. For potential clinical translation, this approach would additionally require in vivo assessments of the therapeutic window for transcriptional up-regulation.

Our CRISPRa experiments revealed another finding of potential future translational relevance. We found that up-regulation of the shorter DSPII isoform was sufficient to rescue cell-cell adhesion function of cardiomyocytes. While DSPII is still too large (6.8 kb) for current AAV vectors, its smaller size could result in more feasibility for gene replacement therapy if vector technology advances.

## Conclusions

This study defines adhesion failure under mechanical load as a core pathogenic mechanism in DSP cardiomyopathy, introduces a valuable platform for investigating desmosome biology, and shows that transcriptional activation of *DSP* is a potential therapeutic strategy.

## Acknowledgements

This work was supported by NIH T32 T32HL007853 (ES), NIH R01-HL149891 and R01-HL105993 (KM), NIH R56-HL157279 (AH), NIH R01-HL171074 (AH), NSF EEC-1647837 (AH and BB). We thank Kevin Ess for his kind gift of the CF-3 control iPSC line. We thank Susan Weintraub for guidance in mass spectrometry experiments.

## Notes

### Competing Interest Statement

The authors have declared no competing interest.

